# Atmospheric hydrogen consumption is regulated by catabolite repression in mycobacteria

**DOI:** 10.1101/2025.11.25.690588

**Authors:** Ashleigh Kropp, James Archer, Marion Jespersen, Thomas Watts, Jessica Solari, Cheng Huang, Ralf B. Schittenhelm, Rhys Grinter, Chris Greening

## Abstract

Consumption of atmospheric hydrogen (H_2_) enables diverse aerobic microorganisms to grow and persist in resource-deprived environments. In the aerobic saprophyte *Mycobacterium smegmatis*, hydrogen oxidation is catalyzed by two differentially expressed, high-affinity, oxygen-insensitive uptake hydrogenases, Huc and Hhy. Huc enables mixotrophic growth and facilitates the transition from growth to dormancy. Although the *huc* operon is known to be upregulated in response to organic carbon deprivation, the specific signals and regulators modulating its expression remain unresolved. Here, we show that GylR, a glycerol-3-phosphate-sensing regulator of glycerol metabolism, plays a role in repression of *huc* expression in response to the availability of glycerol but not other carbon sources. Based on proteomic analyses and activity assays, mutation or knockdown of *gylR* leads to enhanced Huc production and activity. GylR and other key catabolite repressor proteins (Crp1, Crp2) do not directly bind the *huc* operon, indicating repression is mediated by unidentified transcription factors, with GylR acting as an upstream sensor. Here, we present data that suggests atmospheric H_2_ oxidation is regulated in response to organic carbon source availability through the process of catabolite repression. By identifying a key signal that prompts atmospheric H_2_ oxidation, these findings advance understanding of how aerobic bacteria adapt to changing environmental conditions and suggest that organic carbon levels are a key factor regulating the main sink of atmospheric H_2_ in soils globally.

**Importance:** Soil microorganisms collectively consume 70 million tonnes of atmospheric H_2_ a year, regulating atmospheric composition and climate change. In turn, consuming this dependable trace gas enables these microorganisms to survive even when their preferred organic energy sources are exhausted. Despite the importance of H_2_ consumption for soil biodiversity and atmospheric regulation, the signals and sensors that regulate this process remain to be understood. Here, we demonstrate that a model soil bacterium turns on the machinery required for atmospheric H_2_ consumption in direct response to being limited by organic carbon availability, through the process of catabolite repression. Specifically, in the absence of a sensor of the organic carbon source glycerol, a H_2_-consuming hydrogenase is highly expressed and active. These findings suggest that organic carbon levels have a major role in regulating trace gas oxidation, with implications for predicting how trace gas consumption and soil biodiversity respond to environmental change.

## Introduction

Atmospheric trace gas oxidation, through the recently described process of ‘aerotrophy’, provides soil microbes with the metabolic flexibility to grow and persist within resource-deprived environments (1). The oxidation of atmospheric hydrogen is a widespread microbial metabolism, with bacteria and archaea from at least 22 phyla encoding the enzymes capable of catalyzing this reaction, and 9 phyla experimentally confirmed to be atmospheric H_2_ consumers (1, 4, 5). These phyla include many of the dominant bacteria found inhabiting soils (5, 6), making soil communities the primary biogeochemical sink for atmospheric H_2_ and accounting for the net consumption of 75% of H_2_ each year (3, 7). With estimates suggesting that 20-80% of bacteria in any given environment are dormant (1, 8), atmospheric H_2_ oxidation enables microbes to sustain minimal energy requirements while persisting in a non-replicative state, whereby energy demands are substantially reduced (9, 10).

The molecular and cellular basis of atmospheric H_2_ oxidation has been well explored in the aerobic saprophyte *Mycobacterium smegmatis*. This soil actinobacterium can combat resource deprivation by upregulating two high-affinity, oxygen-insensitive [NiFe] hydrogenases, Huc (group 2a) and Hhy (group 1h) (11–13). Huc and Hhy enable *M. smegmatis* to scavenge sub-atmospheric concentrations of H_2_ to support growth and survival (11–13). Differentially expressed, Hhy is active during long-term persistence, whereas Huc enables *M. smegmatis* to grow mixotrophically and is most active during the transition from growth to dormancy (11). The recent structural and functional characterization of Huc has revealed that this enzyme forms an octameric complex composed of the typical large (HucL) and small subunits (HucS), as well as a novel membrane-associated stalk assembled by the medium subunit (HucM). The HucM stalk facilitates long-range menaquinone transport between the hydrogenase active site to the electron transport chain (11, 14) This unique structure supports the capacity of Huc to selectively bind H_2_ and oxidize this substrate to picomolar concentrations, while resisting O_2_ inhibition and remaining stable across a wide range of temperatures (14). The expression of the *huc* operon is tightly regulated, with carbon starvation, oxygen limitation, and elevated H_2_ availability all leading to increased production of this high-affinity hydrogenase (11, 13, 15). Yet the regulatory mechanisms stimulating *huc* expression in response to these conditions remain elusive. For example, it is unresolved whether the induction of Huc during carbon starvation reflects a direct response to organic carbon levels, an indirect response to starvation-induced physiological stress (e.g. redox imbalance), or a general response to the entrance towards stationary phase.

Microorganisms exposed to different environmental and physiological pressures can regulate hydrogenase expression through a variety of mechanisms. Some bacteria capable of growing hydrogenotrophically can directly sense and respond to H_2_ availability. For example, the soil-dwelling proteobacterium *Cupriavidius necator* uses a sensory hydrogenase, to detect environmental H_2_ and upregulate its two key H_2_-consuming hydrogenases through a two-component signal transduction pathway (16–18). Other bacteria upregulate hydrogenases to maintain redox balance and survive amid oxygen deprivation, including *Rhodobacter capsulatus*, which utilises the redox-sensing RegB/RegA two component system to stimulate transcription of a bidirectional hydrogenase when under redox stress (19). Likewise, *M. smegmatis* itself uses the oxygen- and redox-sensing DosST/DosR system to modulate over 50 genes, including those that encode Hhy and a third hydrogenase Hyh that mediates fermentation during hypoxia (13, 15). Organic carbon limitation also drives hydrogenase expression in a variety of organisms. Both *Escherichia coli* and *Salmonella enterica* serovar Typhimurium express uptake hydrogenases during anaerobic conditions when organic carbon availability is limited, a process that is modulated by carbon catabolite repression (20, 21). When the availability of the preferred organic substrate is low, adenylate cyclase produces cyclic AMP (cAMP), which forms a complex with the cAMP receptor protein (CRP) (20, 22). The cAMP-CRP complex then indirectly activates the expression of hydrogenase genes for H_2_ consumption and energy conservation by interacting with unresolved downstream regulators (20–23). Alternatively, when glucose is present in sufficient quantities, cAMP levels are low, and CRP remains unbound, resulting in no activation of hydrogenase expression. This careful modulation of expression ensures that hydrogenase transcription only occurs during organic carbon starvation, meaning that the preferred organic substrate can be metabolized when present without unnecessary investment of resources to make hydrogenases.

Carbon catabolite repression may also play a role in regulating the expression of hydrogenases in *M. smegmatis*, especially given that *huc* and *hhy* are both upregulated in response to organic carbon deprivation (11, 13–15). In *M. smegmatis* and the filamentous soil actinobacterium *Streptomyces coelicolor*, the metabolism of glycerol as an organic carbon source is modulated by the transcriptional regulator GylR, which acts as a cellular sensor of glycerol-derived metabolites (24, 25). In *S. coelicolor*, GylR serves as a direct repressor of the *gylCABX* glycerol metabolism operon in the presence of glycerol, likely through the detection of glycerol-3-phosphate (G3P), and also autoregulates its own expression through a negative feedback loop (25). On the contrary, despite acting as a repressor of the *glpFKD* operon in *M. smegmatis* by directly sensing G3P availability, GylR is also necessary for the activation of genes required for glycerol metabolism (24). In this regulatory pathway, the binding of G3P to GylR induces protein dimerization, resulting in the alleviation transcriptional repression, while simultaneously promoting the recruitment of RNA polymerase for *glpFKD* expression (24). Genomically-encoded regulators known to play distinct roles in catabolite repression, such as GylR and CRP, could potentially modulate hydrogenase expression in *M. smegmatis*; this may either be through directly binding hydrogenase-encoding operons or by functioning as part of a larger regulatory network that represses hydrogenase transcription in the presence of preferred organic substrates.

To address these knowledge gaps, here we performed physiological, biochemical, and proteomic studies to test whether catabolite repression influences *M. smegmatis* hydrogenase production. In addition to informing how environmental factors influence atmospheric H_2_ consumption by soil bacteria, developing a greater understanding of hydrogenase regulation has the potential to inform upscaled production of hydrogenases for industrial purposes, with Huc recently being used to create the first fuel cells powered by atmospheric and waste gas streams (26). We demonstrate that Huc production and activity are regulated in response to glycerol availability through a regulatory network controlled by the glycerol sensor GylR.

## Methods

### Bacterial strains and culture conditions

*M. smegmatis* mc^2^155 and its derivatives (Table S1) were routinely maintained on lysogeny broth (LB) agar plates supplemented with 0.05% (w/v) Tween-80 (LBT) (27). *M. smegmatis* broth cultures were grown in either LBT or Hartmans de Bont (HdB) minimal media (28), each supplemented with 0.05% (w/v) tyloxapol and 0.2% (w/v) of one of four organic carbon sources: glycerol, glucose, acetate or succinate. *E. coli* strains were maintained on LB agar plates and grown in LB broth cultures, unless otherwise specified. For the propagation of pMV261, pLJR962 and pET-23a, media was supplemented with kanamycin (pMV261, pLJR962) (20 µg mL^-1^ for *M. smegmatis* and 50 µg mL^-1^ for *E. coli*) and ampicillin (pET-23a) (100 µg mL^-1^ for *E. coli*). All broth cultures were incubated at 37°C with aeration in a rotary incubation (150-200 rpm). Strains propagated in LBT were inoculated using a single smear of colonies, whilst cultures grown in minimal media were inoculated using turbid LBT cultures, normalised to a starting OD_600_ of 0.01 or 0.06. The consumption of glycerol over time was measured as conducted previously (29). Specific growth rates (µ) were calculated using the follow formula: 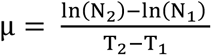, where N_2_ represents the optical density (OD_600_) measured at time T_2_ and N_1_ represents the optical density (OD_600_) measured at time T_1_. Specific growth rates were compared for statistical significance by performing an unpaired t-test with Welch’s correction (p < 0.05) in GraphPad Prism. Timepoints were selected during a period of exponential growth. All bacterial strains and plasmids are listed in Table S1 and S2.

### Isolation and characterization of gylR frameshift mutant

A *gylR* frameshift mutant was spontaneously isolated previously (14). This mutant was analysed using whole-genome sequencing (Peter Doherty Institute, University of Melbourne), and genomic analysis revealed that this strain possessed a single nucleotide insertion at base pair 462 of *MSMEG_*6757, resulting in a frameshift following Leu154 in the transcriptional regulator *gylR*.

### Cloning and molecular biology

Genes encoding proteins for expression or complementation were amplified from *M. smegmatis* mc^2^155 gDNA using primers (Integrated DNA Technologies (IDT)) listed in Table S2. Briefly, genes were amplified by PCR and digested with restriction enzymes (NdeI/XhoI for pET-23a and BamHI/HindIII for pMV261) and ligated into their respective vectors. Vectors were propagated in *E. coli* DH5α before being purified and Sanger sequenced for verification of insert.

### CRISPRi knockdown strain construction

A transcriptional knockdown of *gylR* (*MSMEG_6757*) was constructed as described in Rock *et al*. (30). Briefly, single guide RNA (sgRNA) was designed to the non-template strand of the *MSMEG_6757* gene consisting of 21-bp. Oligonucleotides of this sequence were synthesised by IDT, then annealed and ligated into the kanamycin-selectable CRISPRi plasmid pLJR962 (Addgene plasmid #115162 a gift from Sarah Fortune) using Golden Gate cloning (30, 31) (Table S1). The knockdown construct (pLJR962_KD*gylR*) was transformed with *M. smegmatis* via electroporation and plated onto LBT agar supplemented with 20 mg ml^-1^ kanamycin. Kanamycin-resistant colonies were selected and screened via PCR to confirm genomic integration of pLJR962 contained the desired sgRNA, and knockdown of *gylR* was induced through the addition of 200 ng mL^-1^ anhydrotetracycline (aTc).

### GylR, Crp1 and Crp2 expression and purification

*E. coli* (DE3) C41 transformed with pET-23a(*gylR*), pET-23a(*crp1*) and pET-23a(*crp2*), respectively, were cultured in terrific broth, as previously described (24). Cells were grown at 37°C until OD_600_ = 1.2, followed by induction with 0.3 mM isopropyl-β-d-thiogalactopyranoside (IPTG), and were grown further for 14 hours at 22°C, 180 RPM shaking. Cells were harvested by centrifugation at 5000*g* for 20 minutes, resuspending in Ni-binding buffer (50 mM Tris, 500 mM NaCl, 5% glycerol 20 mM imidazole, pH 8.0 for GylR, 50 mM Tris, 500 mM NaCl, 20 mM Imidazole, pH 7.5 for Crp1 and 50 mM Na_3_PO_4_, 500 mM NaCl, 20 mM Imidazole, pH 7.5 for Crp2) plus 0.1 mg mL^-1^ lysozyme, 0.05 mg mL^-1^ DNase I, and Roche cOmplete^TM^ protease inhibitor cocktail tablet, and lysed by two passages at 40 PSI through cell disruption (Emulsiflex C-5). The resulting lysate was centrifuged at 30,000*g* for 20 minutes and the supernatant applied to HisTrap HP column (Cytiva) i, previously equilibrated in 5 column volumes (CV) binding buffer, followed by washing with 10 X CV of Ni-binding buffer supplemented with 1M NaCl. Proteins were eluted with a step gradient of respective Ni-gradient buffer at of 5%, 10%, 25%, 50% and 100% of 500 mM stock imidazole concentration. For Crp1 and Crp2, fractions containing protein were pooled and concentrated with a 10-kDa molecular weight cutoff (MWCO) concentrator then snap-frozen in liquid N_2_, and stored at −80°C until further use. For GylR, eluted fractions containing the target protein were pooled and applied to a Superdex S200 10/300 SEC column equilibrated in SEC buffer (50 mM Tris pH 8.0, 500 mM NaCl). The respective fractions containing the protein were pooled, concentrated with a 10-kDa molecular weight cutoff (MWCO) concentrator to ∼5 mg mL^-1^ snap-frozen in liquid N_2_, and stored at −80°C until further use. For electrophoretic mobility shift assays, protein was directly used after purification due to instability.

### Huc activity staining

*M. smegmatis* mc^2^155 and its derivatives were cultured in 125 mL or 500 mL conical flasks in 30 mL or 100 mL volumes, respectively, under ambient air conditions. Cultures were harvested at exponential phase (OD_600_ = 1.3 – 1.6 for growth with glycerol, OD_600_ = 1.45 for growth with glucose, OD_600_ = 1.2 for growth with succinate, and OD_600_ = 0.9 for growth with acetate) and stationary phase (OD_max_ + 1 day) via centrifugation (3000*g*, 10 minutes, 4°C), and cell pellets were stored at -20°C. Cell pellets were resuspended in 0.5 mL of lysis buffer (50 mM Tris, 150 mM NaCl pH 8.0), supplemented with 5 mg mL^-1^ of lysozyme, 40 µg mL^-1^ of DNase, 0.25 of Roche cOmplete^TM^ protease inhibitor cocktail tablet. The cell pellet suspension was then lysed using a Constant Systems cell disruptor (40,000 psi, twice), and cell lysate separated from cellular debris via centrifugation (15,000*g*, 10 minutes, 4°C). The protein concentration within each sample was then estimated using a bicinchoninic acid (BCA) assay with bovine serum albumin (BSA) standards. Normalised protein concentrations of each sample were run on pre-cast Native-PAGE 3-12% gels (Invitrogen) or hand-poured Native 7.5% (w/v) BisTris polyacrylamide gels as previously described (32). Pre-cast gels were run at 150 V for 1.5 hours in accordance with the manufacturer’s instructions, whilst hand-poured gels were run in 25 mM Tris 193 mM glycine buffer (pH 8.3) at 25 mA for 3 hours. Gels were run alongside a protein standard (NativeMark Unstrained Protein Standard, Thermo Fisher Scientific), and were visualized using either AcquaStain Protein Gel Stain (Bulldog) or, to assess hydrogenase activity, nitrotetrazolium blue chloride (NBT). For activity staining (14), gels were incubated in 50 mM Tris, 150 mM NaCl pH 8.0 buffer supplemented with 200 µM NBT in an anaerobic Schott bottle amended with a H_2_ anaerobic mix (7% H_2_, 7% CO_2_ in a nitrogen base) for 2 – 24 hours, depending on the level of activity. Activity stains were imaged using a ChemiDoc MP imaging system (Bio-Rad), and hydrogenase activity was determined through the identification of purple-coloured bands of reduced NBT. Where required, the level of Huc activity observed in the Huc oligomer was quantified through densiometric analysis using Image Lab software (Bio-Rad).

### Shotgun proteome analysis

WT, *gylR* mutant and *gylR* knockdown *M. smegmatis* strains were grown in 30 mL volumes, in triplicate, in 125 mL aerated conical flasks containing HdB media supplemented with 0.2% glycerol. Cultures were quenched at exponential phase (OD_600_ ∼1.5) and stationary phase (OD_max_ + 1 day) with 60 mL cold 3:2 glycerol:saline solution (−20°C). Cultures were subsequently harvested by centrifugation (4,800g, 30 min, −9°C), further quenched with 1 mL cold 1:1 glycerol:saline solution (stored at −20 °C), and pelleted before washing in ice-cold phosphate-buffered saline (PBS). To lyse the cell pellets and denature proteins, the pellets were resuspended in lysis buffer (50 mM Tris-HCl, pH 8.0, 2 mM MgCl_2_, lysozyme, DNase) supplemented with sodium dodecyl sulfate (SDS) (final concentration of 4%). Samples were boiled at 95 °C for 10 minutes and sonicated (Bioruptor, Diagenode) using 20 cycles of ‘30 seconds on’ followed by ‘30 seconds off’, remaining on ice in-between cycles. The lysates were clarified by centrifugation (14,000 × g, 10 minutes, room temperature). Protein concentration was confirmed using the BCA assay kit (Thermo Fisher Scientific) and equal amounts of protein were processed from the strains in exponential and stationary phase for downstream analyses. After removal of SDS by chloroform/methanol precipitation, the proteins were proteolytically digested with trypsin (Promega) and purified using OMIX C18 Mini-Bed tips (Agilent Technologies) prior to LC-MS/MS analysis. Using a Dionex UltiMate 3000 RSLCnano system equipped with a Dionex UltiMate 3000 RS autosampler, the samples were loaded via an Acclaim PepMap 100 trap column (100 µm × 2 cm, nanoViper, C18, 5 µm, 100 Å; Thermo Scientific) onto an Acclaim PepMap RSLC analytical column (75 µm × 50 cm, nanoViper, C18, 2 µm, 100 Å; Thermo Scientific). The peptides were separated by increasing concentrations of (80% acetonitrile/0.1% formic acid) for 158 minutes and analysed with an Orbitrap Fusion Tribrid mass spectrometer (Thermo Scientific) operated in data-dependent acquisition mode using in-house, LFQ-optimized parameters. Acquired .raw files were analysed with MaxQuant to globally identify and quantify proteins across conditions (33). Data visualization and statistical analyses were performed in Perseus (34).

### Electrophoretic mobility shift assay

The binding of GylR to the *huc* promoter was investigated through electrophoretic mobility shift assays using DIG Gel Shift Kit, 2nd Generation (Roche). A 454-bp *huc* promoter (P*_huc_*) was amplified using primers hucp_fw and hucp_rev (13). In addition, a 306-bp *glpFKD* promoter (P*_glpFKD_*) was amplified using glpp_fw and glpp_rev primers (24). The amplified products were purified, concentrated, and labelled with digoxigenin (DIG) at the 5’ end according to the manufacturer’s protocol (Roche). Next, DNA-protein reactions (20 µl) were prepared containing 0 ng, 25 ng, or 75 ng of purified GylR, Crp1 or Crp2 protein and 310 fmol of DIG-labelled P*_huc_* or P*_glpFKD_* in binding buffer (20 mM HEPES, pH 7.6, 1 mM EDTA, 10 mM (NH_4_)_2_SO_4_, 1 mM DL-Dithiothreitol (DTT), 0.2% (w/v) Tween 20, 30 mM KCl). Selective reactions also contained 50 mM of glycerol-3-phosphate (G3P) or cAMP. The reaction mixtures were incubated for 15 minutes at room temperature. Next, a 5% non-denaturing polyacrylamide gel, prepared as per manufacturer’s protocol (Roche), was pre-run for 60 min at 6-18 mA in 0.5 × TBE buffer (44.5 mM Tris, 44.5 mM boric acid, 1 mM EDTA, pH 8.0). After pre-run, the DNA-protein reaction mixtures were loaded onto the gel and was run at 6-15 mA until the dye front was two-thirds down the gel, followed by contact transfer to a NYLM-RO nylon membrane (Roche), as described by manufacturer’s protocol (Roche). DIG-labelled free DNA and DNA–protein complexes were detected according to the manufacturer’s protocol (Roche).

## Results and Discussion

### GylR modulates Huc activity in response to glycerol availability

To investigate the importance of GylR for *M. smegmatis* growth, wild-type (WT) *M. smegmatis* and a *gylR* frameshift mutant strain were grown in minimal media supplemented with glycerol as the sole carbon source (14). The growth rate of *gylR* mutant was much slower compared to WT *M. smegmatis* (Figure 1A), though the cells reached the same growth yield, suggesting the strain uses glycerol less rapidly. This is consistent with previous observations that GylR is a transcriptional activator of the *glpFKD* operon (24), with the absence of a functional *gylR* reducing the expression of genes required for glycerol import (glycerol uptake facilitator) and consumption (glycerol kinase, glycerol-3-phosphate dehydrogenase) needed for rapid growth on glycerol. The *gylR* mutant strain was still able to consume glycerol despite the inactivation of GylR, albeit at a slower rate (Figure 1B); this suggests that the *glpFKD* operon is only partially repressed in the *gylR* mutant strain, consistent with previous literature demonstrating that the glycerol metabolism operon is under the control of other transcriptional regulators, such as Crp and SigF (24). The observed growth defect in the *gylR* mutant strain was solely due to the inactivation of GylR, with complementation restoring growth to the WT phenotype (Figure 1A).

**Figure 1:**
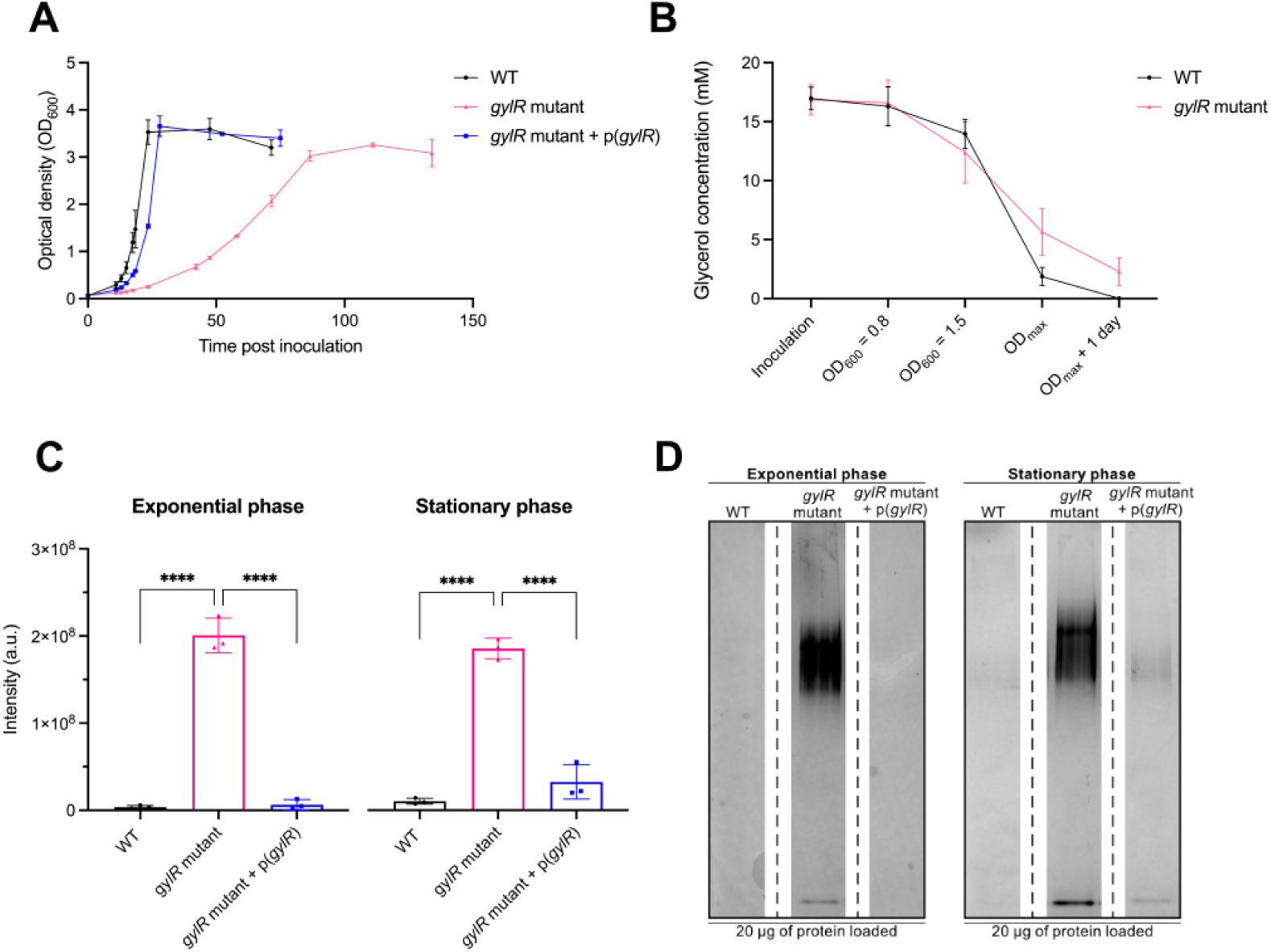
The inability to efficiently utilize glycerol increases Huc activity when *M. smegmatis* is grown with glycerol as the sole carbon source. **(A)** Cell density time-course demonstrating the comparative growth of WT *M. smegmatis* (WT + pMV261(empty), *gylR* mutant (*gylR* mutant + pMV261(empty)) and *gylR* mutant with *gylR* complementation (*gylR* mutant:p(*gylR*)). **(B)** Glycerol consumption monitored at different growth phases in both WT *M. smegmatis* and the *gylR* mutant frameshift mutant. **(C)** Quantification of Huc activity in WT *M. smegmatis* (WT + pMV261(empty), *gylR* mutant (*gylR* mutant + pMV261(empty)) and *gylR* mutant with *gylR* complementation (*gylR* mutant:p(*gylR*)) through densitometric analysis of native gels (Figure S1). Strains were grown as three independent biological replicates (n = 3) with glycerol as the sole carbon source, and harvested at exponential phase (OD_600_ = 1.4-1.6) and stationary phase (OD_max_ + 1 day). The intensity of the reduced artificial electron acceptor nitrotetrazolium blue chloride (NBT) present on the gels was quantified using Image Lab. The level of activity between strains at exponential and stationary phase was respectively compared for statistically significant differences using a one-way ANOVA with Tuckey’s multiple comparisons test (p < 0.05), with error bars demonstrating the standard deviations of the three biological replicates. **(D)** Native-PAGE hydrogenase activity staining of WT *M. smegmatis* (WT + pMV261(empty), *gylR* mutant (*gylR* mutant + pMV261(empty)) and *gylR* mutant with *gylR* complementation (*gylR* mutant:p(*gylR*)) using the artificial electron acceptor NBT. Cells were harvested at in triplicate (n=3) at exponential phase (OD_600_ = 1.4-1.6) and stationary phase (OD_max_ + 1 day). 20 µg of each sample was loaded, and only one replicate for each strain/growth stage is illustrated. Upper red arrow indicates oligomeric Huc staining and lower red arrow indicates dimeric Huc staining.

Because the *gylR* mutant grows slowly on glycerol, we hypothesised that enzymes like Huc that support persistence or alternative energy capture might be upregulated. To establish whether GylR plays a role in regulating the expression of the *huc* operon, we assayed Huc activity in the *gylR* mutant and WT *M. smegmatis* strains when grown with glycerol as the only carbon source. Activity staining of Huc was performed using whole-cell lysates of WT and *gylR* mutant strains, grown to either exponential phase (OD_600_ = 1.4-1.6) or stationary phase (OD_max_ + 1 day), and the activity of the Huc was quantified using densitometry (Figure 1D upper red arrow; Figure S1) (14). As observed in previous studies (11, 35), the activity of Huc was minimal at exponential phase in the WT strain (average intensity 3.72 × 10^6^ absorbance units (a.u.)) (Figure 1C and 1D), most likely due to the presence of glycerol for mixotrophic growth (Figure 1B). WT cells exhibited the highest level of Huc activity at stationary phase (average intensity 1.03 × 10^7^ a.u.), where *M. smegmatis* cells are transitioning from growth to dormancy due to the onset of carbon starvation and depletion of glycerol (Figure 1B, 1C and 1D). Comparatively, the *gylR* mutant strain exhibited significantly higher Huc activity compared to the WT strain, with staining of the Huc oligomer exhibiting an average intensity of 2.01 × 10^8^ and 1.85 × 10^8^ a.u. at exponential and stationary phase respectively (Figure 1C and 1D). Notably, the level of Huc activity at exponential phase was approximately 54-fold greater in the *gylR* mutant strain compared to WT *M. smegmatis*. The increase in Huc activity in the *gylR* mutant strain was exclusively due to the presence of a non-functional GylR, with complementation of the *gylR* mutant strain restoring Huc activity to WT levels (Figure 1C and 1D; Figure S1).

### GylR is not required for growth and Huc activity with alternative organic substrates

Next, we aimed to investigate whether the modulation of Huc activity by GylR was influenced by the availability of alternative organic substrates. To this end, WT and *gylR* mutant *M. smegmatis* strains were grown in minimal media supplemented with one of four organic substrates: glycerol, glucose, acetate or succinate, as the sole carbon source. When grown with glycerol, a statistically significant reduction (p < 0.05) in the growth rate of the *gylR* mutant (µ = 0.0379 ± 0.0037 h^-1^) was observed compared to WT *M. smegmatis* (µ = 0.1971 ± 0.0.002 h^-1^) (Figure 2A; Figure S2). In contrast, the growth rate of the *gylR* mutant with glucose, acetate or succinate was not significantly different from WT (Figure 2A and S2). This difference in the growth of the *gylR* mutant between carbon sources highlights the importance of GylR for glycerol metabolism, confirming that this regulator does not modulate the expression of genes required for the metabolism of alternative organic substrates.

**Figure 2:**
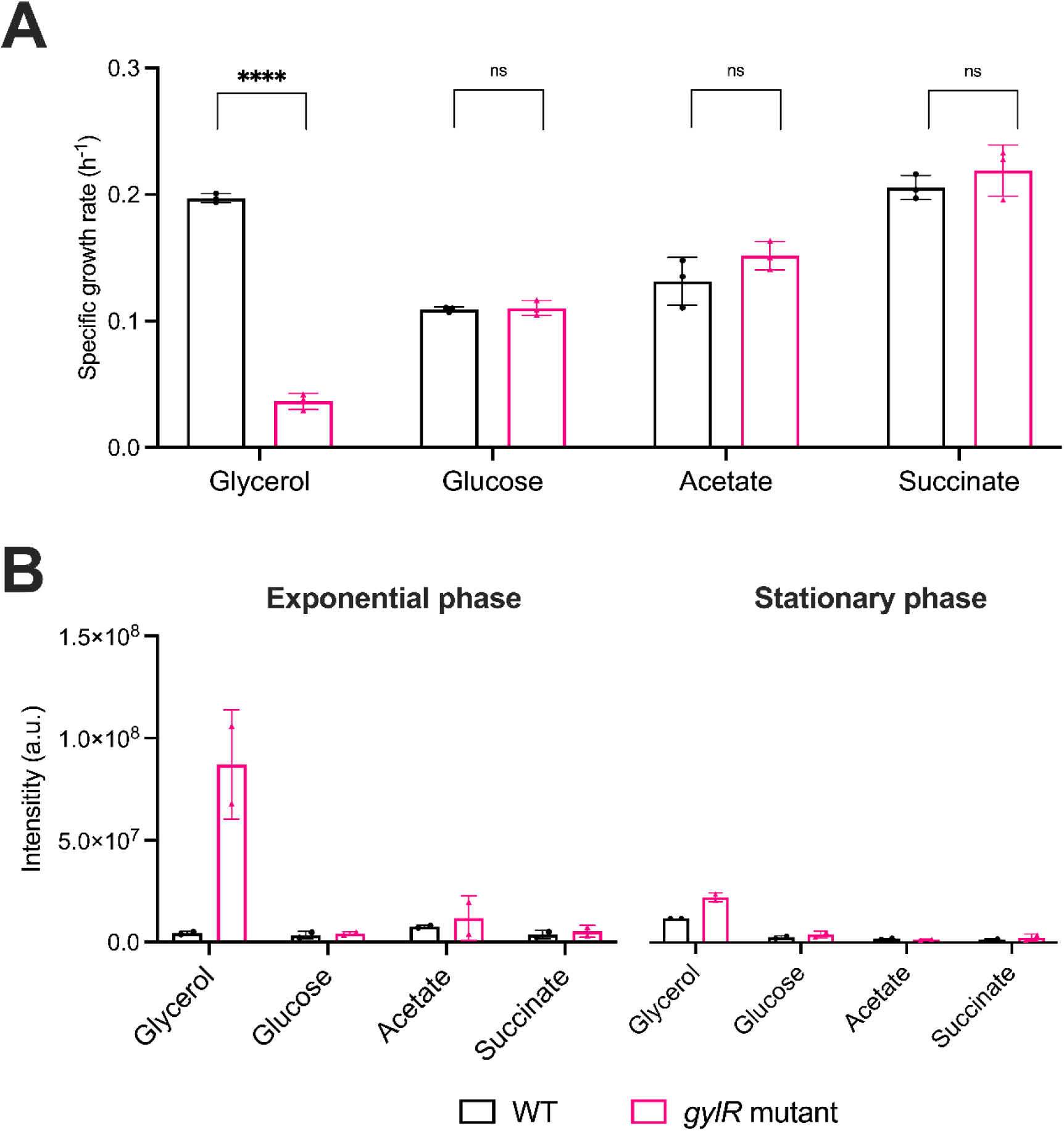
A lack GylR does not impact *M. smegmatis* growth or Huc activity on carbon sources other than glycerol. **(A)** Comparison of the specific growth rates of WT and the *gylR* mutant strains grown in minimal media supplemented with glycerol, glucose, acetate or succinate as the sole carbon source. Statistical comparison of specific growth rates was performed using an unpaired t-test with Welch’s correction (p < 0.05), with error bars demonstrating the standard deviations of three (n=3) biological replicates. **(B)** Quantification of Huc activity in WT *M. smegmatis* and the *gylR* frameshift mutant through densitometric analysis of native gels (Figure S3). Strains were grown as two independent biological replicates (n = 2) with glycerol, glucose, acetate or succinate as the sole carbon source. Strains were harvested at exponential phase (OD_600_ = 1.3 – 1.5 for growth with glycerol, OD_600_ = 1.45 for growth with glucose, OD_600_ = 1.2 for growth with succinate, and OD_600_ = 0.9 for growth with acetate) and stationary phase (OD_max_ + 1 day). The intensity of the reduced artificial electron acceptor NBT present on the gels was quantified using Biorad Image Lab.

We next investigated whether Huc activity was regulated by GylR in response to the availability of different carbon sources in *M. smegmatis*. During growth with glucose, acetate or succinate as the sole carbon source, no difference in Huc activity was observed between the WT and *gylR* mutant strains, in contrast to the substantially increased Huc activity observed by the *gylR* mutant during growth with glycerol (Figure 2B; Figure S3). Altogether, the increase of active Huc in the absence of a functional GylR strongly suggests that this regulator is involved in the modulation of *huc* expression in response to glycerol availability, in addition to regulating glycerol transport and consumption genes. Importantly, the increase in Huc activity across both phases suggests this is the result of GylR playing a role in the repression of *huc* expression when glycerol is present.

### M. smegmatis increases Huc production when unable to sense glycerol availability

Using untargeted quantitative shotgun proteomics, we confirmed that increased Huc activity in the *gylR* mutant strain is due to increased production of the Huc enzyme. Substantial proteome differences were observed between the *gylR* mutant and WT *M. smegmatis* strains grown in minimal media with glycerol as the sole carbon source, with 396 and 555 proteins differing significantly in abundance during exponential and stationary phases, respectively (Figure 3A and B). During exponential phase, all structural subunits of Huc (HucL, HucM and HucS) were significantly more abundant in the *gylR* mutant compared to WT, increasing 16-, 22- and 203-fold, respectively (Figure 3A). The abundance of the structural subunits also increased significantly at stationary phase, by 9.7-, 14- and 11-fold, respectively (Figure 3B). Moreover, the *huc*-associated auxiliary proteins HypC-E, which mature the active site of [NiFe] hydrogenases, increased in abundance in the *gylR* mutant strain, highlighting their requirement for rapid Huc assembly (Figure 3A and 3B) (36, 37). As expected, the products of the *glpFKD* operon were also less abundant in the *gylR* mutant strain, with glycerol kinase (GlpK) and glycerol-3-phosphate dehydrogenase (GlpD) decreasing 267- and 792-fold at exponential phase, respectively (Figure 3A) (24), whereas no differential abundance was detected for the glycerol uptake facilitator (GlpF). Proteome analysis of a CRISPRi knockdown of *gylR* in comparison to WT further validated the phenotype exhibited by the *gylR* mutant strain, with the transcriptional silencing of this gene resulting in the elevated abundance of the three primary structural subunits of Huc, as well as auxiliary proteins required for [NiFe] hydrogenase assembly, at both exponential and stationary phase (Figure S4).

**Figure 3:**
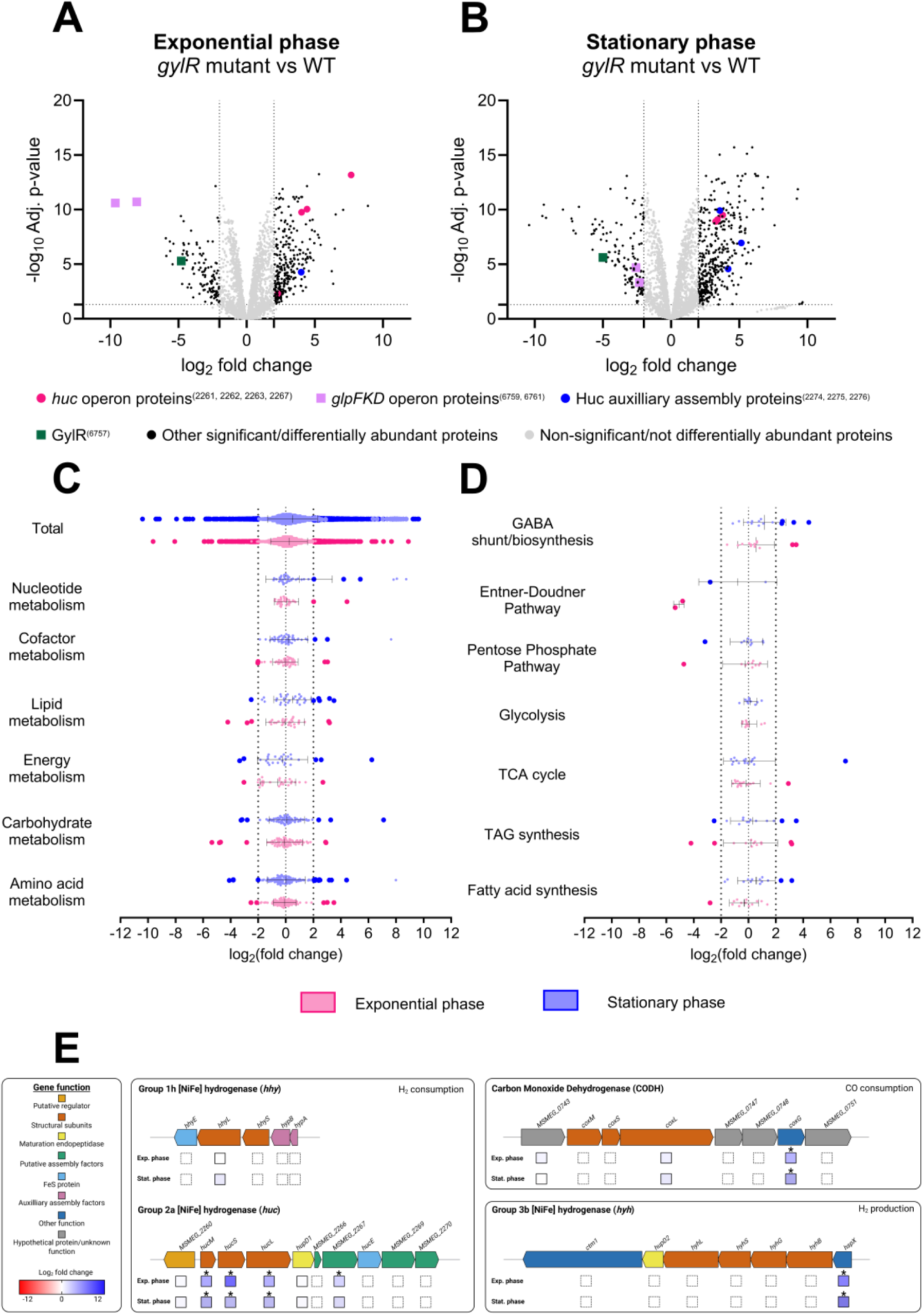
Huc proteins are significantly more abundant when GylR is inactive. Comparative shotgun proteomics volcano plots demonstrating the differential abundance of proteins in the *gylR* mutant strain compared to WT *M. smegmatis* when grown with glycerol as the sole carbon source at exponential phase (OD_600_ = 1.5) **(A)** and carbon-depleted stationary phase (OD_max_ + 1 day) **(B)**. log_2_fold change represents the ratio of abundance in the *gylR* mutant strain vs WT *M. smegmatis* (n=3), using an adjusted p-value threshold of 0.05. Global changes in the proteome of the *gylR* mutant vs WT *M. smegmatis* were examined by assigning proteins with predictive metabolic functions and were categorised based on KEGG pathways **(C)** and KEGG modules **(D)** at exponential phase (pink) and stationary phase (blue). Proteins with statistically significant differences in abundance (p < 0.05, log_2_ FC ≥ 2, log_2_ FC ≤ -2) are represented by the large dark-coloured dots. All error bars represent standard deviation from the mean. **(E)** Genetic organisation of the three [NiFe] hydrogenases and carbon monoxide dehydrogenase (CODH) of *M. smegmatis*. The log2 fold change of proteins encoded by each gene is represented by a colour-scaled box for both exponential and stationary phase. Boxes with an asterisk (*) indicate that the fold-change is statistically significant (p < 0.05) (Table S3). Boxes with a dashed outline indicate that the relevant protein was not detected in the dataset. Figure 3E was made using BioRender.

Collectively, the striking increase in Huc activity and elevated abundance of *huc*-encoded proteins observed in the *gylR* frameshift mutant highlights the role of GylR in the repression of *huc* when glycerol is available as the preferred organic substrate (Figure 1C, Figure 3A and 3B). Although *huc* expression is typically upregulated during the transition from growth to dormancy (11), the strong induction of Huc activity and production at exponential phase in the *gylR* mutant, whereby glycerol is still available for growth, is consistent with *huc* being regulated by catabolite repression. The structural subunits of other trace gas oxidizing enzymes, namely for the Hhy hydrogenase (HhyLS) and carbon monoxide dehydrogenase (CoxLMS), exhibit no significant change in abundance in the proteomic dataset (Figure 3A, 3B and 3E) (11, 13, 29, 38). With these enzymes typically upregulated following organic carbon starvation later into persistence, Huc production could be triggered initially by the absence of the glycerol catabolite G3P. This precise modulation of hydrogenase expression, with the *huc* operon repressed in the presence of the preferred energy source glycerol and induced during glycerol limitation, reduces unnecessary energy expenditure of *M. smegmatis* during growth under nutrient-replete conditions

Further system-wide analysis of the proteomic data provided insights into how *M. smegmatis* remodels its metabolism in response to glycerol availability. We first compared the proteomic changes in the *gylR* mutant at exponential and stationary phase with previously published gene-expression data from *M. smegmatis* grown in continuous culture with glycerol as the sole carbon source (39). We observed a significant correlation between proteins that changed in abundance in the *gylR* mutant relative to wild type, and genes differentially expressed under slow versus fast dilution rates (Figure S5). This correlation was stronger in the *gylR* mutant exponential-phase proteome than in the stationary-phase proteome, suggesting that *M. smegmatis* responds similarly to low environmental glycerol availability and to the loss of GylR-mediated transcriptional regulation (Figure S5). This likely reflects that a large proportion of genes differentially expressed during glycerol starvation are regulated primarily by sensing of glycerol availability *via* GylR, whereas other genes are induced through independent mechanisms in response to other external and physiological signals. Indirectly, the moderately decreased growth rate and substrate consumption of the *gylR* mutant may also induce a mild starvation response that modulates gene expression. However, the Huc production and activity observed in the *gylR* mutant background is unprecedentedly high and greatly exceeding for example the induction seen during starvation-induced stationary phase in the wild-type background. This induction primarily reflects the direct effects of loss of catabolite repression, whereas indirect effects would only induce marginal additional changes.

Proteins associated with key glycerol metabolism pathways, such as the Entner-Doudoroff Pathway (EDP) and Pentose Phosphate Pathway (PPP), were significantly less abundant in the *gylR* mutant compared to WT *M. smegmatis*, highlighting the inability of the *gylR* mutant strain to efficiently sense glycerol and initiate the expression of genes essential for its catabolism (Figure 3C and Figure 3D;Table S3). As a result, carbon flux appeared to be redirected to alternative pathways, with key enzymes for triacylglycerol (TAG) and gamma-aminobutyric acid (GABA) synthesis significantly more abundant in the *gylR* mutant compared to WT *M. smegmatis* (Figure 3D;Table S3). The synthesis of TAGs has been previously observed as a preparatory mechanism for dormancy in *M. smegmatis*, providing an energy reservein the form of fatty acids, whilst the GABA shunt pathway supplies the tricarboxylic acid (TCA) cycle with succinate from glutamate, rather directly from α-ketoglutarate in the typical TCA cycle (40–45). The divergence of carbon flux through these pathways was further supported by the increased abundance of the flavoprotein subunit of the succinate dehydrogenase 1 (Sdh1) (Figure 3D;Table S3). This enzyme is non-essential for *M. smegmatis* growth, but crucial for the oxidation of succinate to fumarate, delivering electrons to the respiratory transport chain to drive ATP synthesis (46, 47). Collectively, the use of alternative carbon catabolism pathways likely reflects a coordinated metabolic response to glycerol availability, with GABA and TAG synthesis providing important intermediates for the TCA cycle that enables the production of ATP and assists in maintaining redox homeostasis (40–45, 48). Moreover, the metabolic remodeling observed in the *gylR* mutant strain emphasizes the role of GylR as the primary glycerol sensor for *M. smegmatis*, with the absence of a functional sensor preventing the induction of glycerol metabolism through high-yielding energy pathways, such as EDP and PPP.

### GylR does not directly repress transcription of the huc operon

To investigate whether GylR represses *huc* expression directly, we examined whether GylR binds to the promoter region upstream of the *huc* operon. Electrophoretic Mobility Shift Assays (EMSA) were performed using recombinantly purified GylR (Figure S6), which was incubated with the previously confirmed *huc* promoter region (13). We did not observe GylR interaction with the *huc* promoter, irrespective of the concentration of protein incubated with the promoter DNA (Figure 4A). The addition of G3P, the catabolite of glycerol, which interacts with GylR to modulate the expression of the *glpFKD* operon in response to glycerol availability (24), had no impact on the binding of GylR to the *huc* promoter (Figure 4A). As expected, the presence of GylR at increasing concentrations resulted in a shift of the *glpFKD* promoter DNA, with the addition of G3P reducing this shift (Figure 4B and 4C, red arrows). The lack of GylR binding to the *huc* promoter indicates that GylR does not directly repress *huc* expression. Instead, GylR may repress an activator of *huc,* resulting in low expression in the presence of G3P. This result further supports the idea that *huc* is regulated by catabolite repression, specifically in response to glycerol availability, through a signal transduction cascade initiated by the sensor GylR.

**Figure 4.**
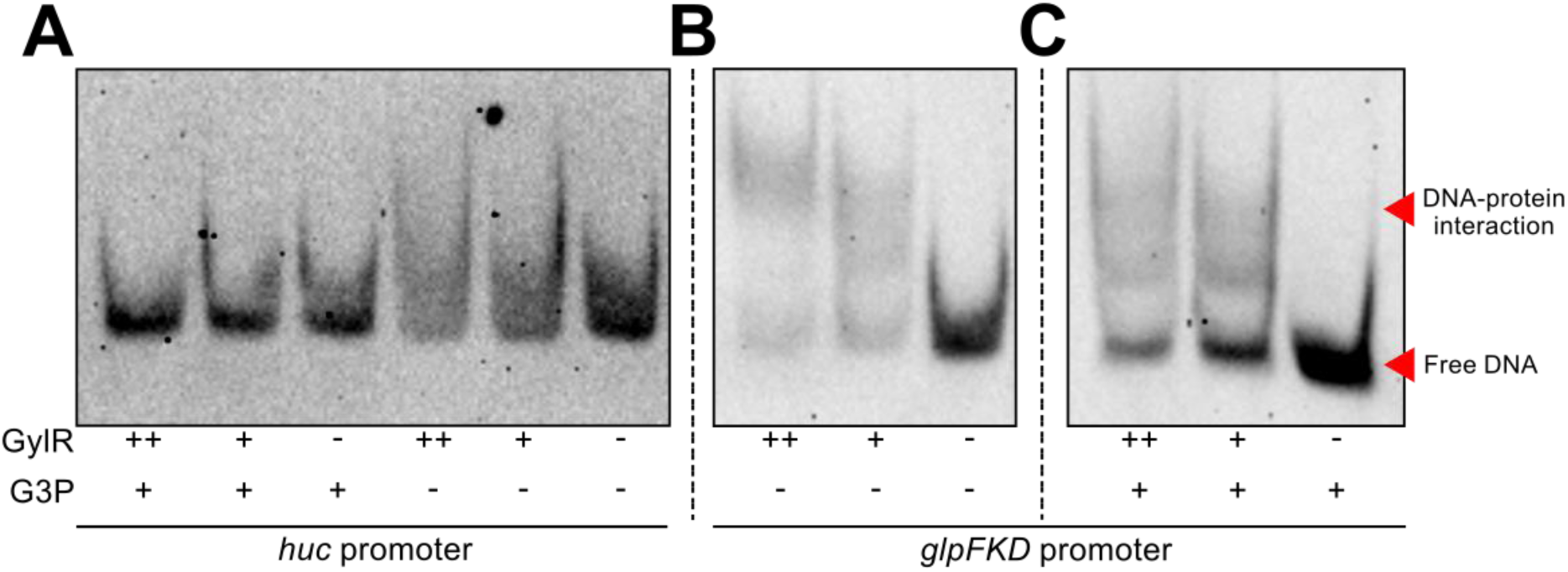
GylR does not directly bind to the huc promoter. **(A)** EMSA assessing potential binding of purified GylR to the *huc* operon, with and without G3P, an effector molecule reported to modulate binding of GylR to the *glpFKD* promoter (24). The binding of GylR to the *glpFKD* promoter was used as a positive control (24), with the absence **(B)** and addition **(C)** of G3P. DNA-protein interaction is indicated by the upward shift in band molecular weight as protein concentration increases. Upper red arrow indicates gel shift and lower red arrow is free DNA.

We hypothesised that alternatively CRP may be a transcription factor that directly regulates Huc expression, given it is a major transcriptional regulator in *M. smegmatis* and a mediator of catabolite repression of hydrogenases in Enterobacteriaceae (20–23, 49–52). In *M. smegmatis*, two CRP homologues, Crp1 (*MSMEG_0539*) and Crp2 (*MSMEG_6189*), regulate mycobacterial metabolism, including the *glp* operon (24). Mobility shift assays using purified Crp1 or Crp2 (Figure S6) indicated that neither protein binds to the *huc* promoter in the presence or absence of cAMP (Figure S7) (52). Additionally, hypothetical transcriptional regulators identified from a DNA pull-down using the *huc* promoter region (MSMEG_3822, MSMEG_2386, MSMEG_0916, MSMEG_2600) were also investigated through mobility shift assays; however, these proteins were found not to bind the *huc* promoter and were likely a result of indirect association with the DNA (data not shown). Collectively, these data suggest the presence of an unidentified regulators downstream of GylR that modulates *huc* expression, with future work needed to unravel this regulatory network.

## Conclusions

Here we provide the first demonstration that the ecologically and biogeochemically critical process of atmospheric trace gas oxidation is directly regulated by organic carbon availability. Although atmospheric H_2_ oxidation is well known to be induced by carbon starvation in *M. smegmatis* and other bacteria, it has remained unclear whether this response reflects an environmental signal (i.e., organic carbon levels), a physiological cue (e.g., redox or electron imbalance), or a broader transcriptional program associated with entry into stationary phase. We found that in response to a non-functional GylR resulting in the inability to sense glycerol-3-phosphate and rapidly metabolise glycerol, Huc is overexpressed and highly active. We propose that GylR is an indirect repressor of Huc expression when glycerol is abundant, demonstrating that catabolite repression controls Huc expression in *M. smegmatis*. This form of catabolite repression ensures that H_2_ oxidation is activated only as a metabolic last resort, enabling cells to prioritise rapid growth when organic carbon is abundant and to shift toward atmospheric energy scavenging during scarcity. Such regulation mirrors the logic of catabolite repression in *Enterobacteriaceae* but is adapted in *Mycobacterium* to sustain survival by exploiting a universally available atmospheric energy source. Key questions nevertheless remain: What are the direct transcription factors controlling Huc repression that act downstream of GylR? Do other catabolite sensors regulate Huc in the presence of other organic carbon sources? And how are Hhy and CO dehydrogenase induced during starvation, given they are unresponsive to GylR? A combination of targeted (e.g. promoter pulldowns) and untargeted (e.g. transposon mutagenesis screens) approaches may help to identify further regulators, though the former approach did not yield any specific *huc*-binding proteins.

These findings have broad implications and applications. From a biotechnology perspective, by relieving its repression, we have been able to produce sufficient Huc to generate the first air-powered fuel cells (14, 26, 53, 54). More broadly, this study also improves the understanding of how the biological sink of atmospheric H_2_ is regulated. Organic carbon is one of the most important environmental factors predicting the abundance, expression, and activity of atmospheric H_2_ oxidation in the environment (1, 3–5, 12, 55–58). This likely reflects that Actinobacteriota (including *Mycobacterium*) are the dominant sinks of atmospheric H_2_ globally and many likely adopt analogous catabolite repression mechanisms to adapt to resource variability and limitation (1, 4, 5, 7, 12, 55, 57–59). As such, changes in soil carbon availability due to various factors (e.g. land use change, fertilisation, warming) may directly influence the strength of the global H_2_ sink by modulating the expression and activity of high-affinity hydrogenases across microbial communities.

## Contributions

C.G, A.K and R.G conceived, designed and supervised the study. Different authors were responsible for culture preparation and harvesting (A.K, J.A, M.J, T.W, J.S), growth analysis (A.K and J.A), glycerol consumption analysis (J.S), activity staining (J.A, M.J and A.K), shotgun proteomic analysis (A.K, J.A, T.W, C.H and R.B.S), molecular biology (A.K), protein purification (A.K) and EMSAs (A.K). J.A, A.K and C.G wrote the manuscript with input from all authors.

## Acknowledgements

This work was supported by ARC Discovery Project grants (DP200103074, DP230103080; to C.G. and R.G.), an Australian Government Research Training Program Stipend (to A.K. and J.A.), and an NHMRC EL2 Fellowship (APP1178715; salary for C.G.) and ARC Future Fellowship (FT240100502; salary for C.G.). We thank Dr. George Taiaroa and A/Prof Debbie Williamson for sequencing the mutant. Paul R. F. Cordero contributed to *gylR* identification, and Thanavit Jirapanjawat and Luis Jimenez provided technical assistance.

