## Supplementary material for "Atmospheric hydrogen consumption is regulated by catabolite repression in mycobacteria"

**Table S1. Bacterial strains and plasmids used in this study.**

| **Strain** | **Description** | **Reference** |
|---|---|---|
| $mc^2155$ | Wildtype strain of *Mycobacterium smegmatis* | (60) |
| *gylR* mutant | Isogenic strain to WT, but possesses a frameshift mutation (Leu154) to *MSMEG_6757* (*gylR*) | This study |
| WT + pMV261 | WT strain with empty pMV261 for complementation assay | This study |
| *gylR* mutant + pMV261 | *gylR* mutant strain with empty pMV261 for complementation assay | This study |
| *gylR* mutant + p(*gylR*) | *gylR* mutant strain with pMV261(*gylR*) for complementation assay | This study |
| *gylR* knockdown | WT strain with pLJR962_KDgylR | This study |
| DH5α | *Escherichia coli* F– ϕ80*lacZ*ΔM15 Δ(*lacZ*YA*argF*)U169 *recA*1 *endA*1 *hsdR*17(rK–, mK+) *phoA supE*44 λ– *thi*-1 *gyrA*96 *relA*1 | Thermo Fischer |
| C41 (DE3) | Standard *Escherichia coli* lab strain used for recombinant protein expression | Thermo Fischer |
| **Plasmids** | **Description** | **Reference** |
| pMV261 | $Kan^r$, mycobacterial oriM, pBR322 ori, $P_{hsp60}$ promoter | (61) |
| p(*gylR*) | pMV261 containing *MSMEG_6757* (*gylR*) gene insert for *gylR* mutant complementation | This study |
| pET-23a | Protein expression vector possessing N-terminal T7-Tag and $Amp^r$ | Novagen |
| pET-23a(*gylR*) | pET-23a containing *gylR* sequence for recombinant protein expression. | This study |
| pET-23a(*crp1*) | pET-23a containing *crp1* sequence for recombinant protein expression. | This study |

| | | |
|---|---|---|
| pET-23a(*crp2*) | pET-23a containing *crp2* sequence for recombinant protein expression. | This study |
| pLJR962 | Sth1 dCas9; Sth1 sgRNA scaffold; Tet repressor; L5- integrating backbone; ColE1 ori (E. coli); $Kan^r$ | (30) |
| pLJR962_KDgylR | pLJR962 with sgRNA targeting *gylR* for repression | This study |

**Table S2. Primers used in this study.**

| **Primers** | **Sequence (5' to 3')** | **Purpose** |
|---|---|---|
| hucp_fw | CGACCAGACGCGCGGCCTC | Amplification of *huc* promoter of EMSA |
| hucp_rev | GACCGGCGAGATGTCTGGAAGTTC | |
| glpp_fw | GTAGCTGCAGTATCGCCGCGG | Amplification of *glpFKD* promoter of EMSA |
| glpp_rev | CACCCCGAACAGGATGAGGATGC | |
| gylrC_fw | GTCAGGGATCCATGCCAGGCACTGTGCAGTCCGTG | Amplification of *gylR* for complementation vector construction |
| gylrC_rev | GTCAGAAGCTTTCACAGTTCCCGCCCGTGGC | |
| gylR_fw | GTCAGCCATGGGTCCAGGCACTGTGCAGTCCGTG | Amplification of *gylR* for recombinant protein expression |
| gylR_rev | GTCAGCTCGAGCAGTTCCCGCCCGTGGCC | |
| crp1_fw | GTCAGCCATGGGTGACGAAGTGCTGGCGCGCG | Amplification of *crp1* for recombinant protein expression |
| crp1_rev | GTCAGAAGCTTTCAGTTCGCGCGCCGCGC | |
| crp2_fw | GTCAGCCATGGGTGACGAGATCCTGGCCAGGGC | Amplification of *crp2* for recombinant protein expression |
| crp2_rev | GTCAGAAGCTTCTAGCGGGCGCGGCGGGC | |
| gylr_KD_fw | GGGAATCCAGTTGGGTCGAACCGAG | sgRNA targeting *gylR* repression |
| gylr_KD_rev | AAACCTCGGTTCGACCCAACTGGAT | |

| | | |
|---|---|---|
| pljr962_fwd | GCTCTTCAGGATCTGACCAGGGAAAATAGCCCTC | Screening of strains with pLJR962_KDgylR |
| pljr962_rev | GCTCTTCACTGAAAAAAATAAAAAAGGGGACCTCTA | |

**Table S3 (xlsx). Summary of proteomic analysis data.** Spreadsheet details the results of the shotgun proteomic experiment, which compared the relative abundance of proteins in the gylR mutant strain vs WT *M. smegmatis*, as well as a *gylR* knockdown vs WT *M. smegmatis*, at exponential phase and stationary phase. The spreadsheet also contains known *M. smegmatis* proteins mapped to KEGG pathways and modules, with annotations assigned to proteins identified in the *gylR* mutant and WT strains.

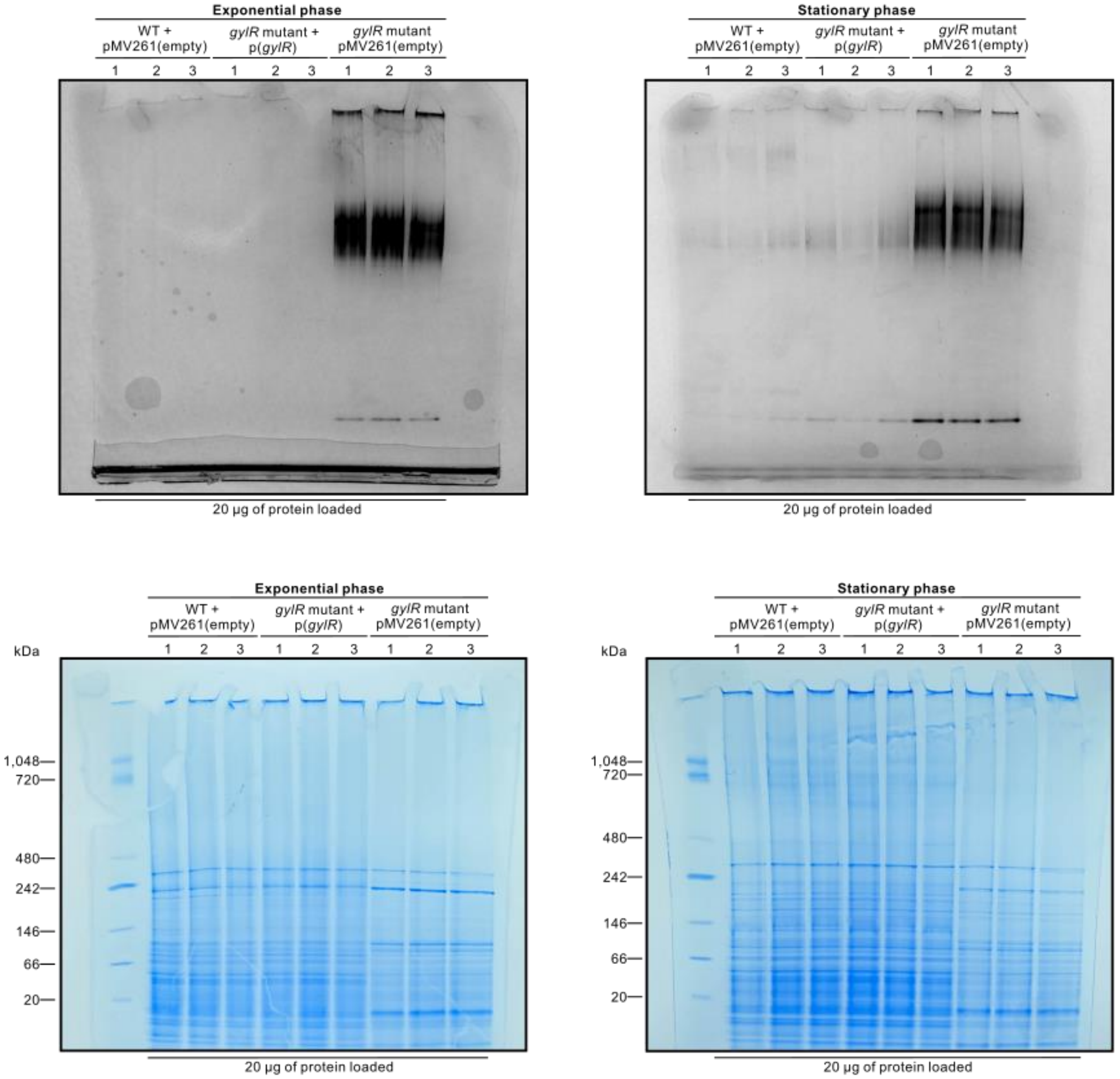


**Figure S1. Native-PAGE hydrogenase activity staining of the *gylR* mutant frameshift mutant**. Native gels depicting Huc hydrogenase activity stained with the artificial electron acceptor NBT (top) and Coomassie gel stained using AcquaStain (bottom). WT *M. smegmatis* (WT +

pMV261(empty), *gylR* mutant (*gylR* mutant + pMV261(empty) and the *gylR* mutant complementation strain (*gylR* mutant:p(*gylR*)) were harvested in triplicate (n=3) at exponential phase ($OD_{600}$ = 1.4-1.6) and stationary phase ($OD_{max}$ + 1 day) for activity quantification using densitometry. 20 µg of each sample was loaded.

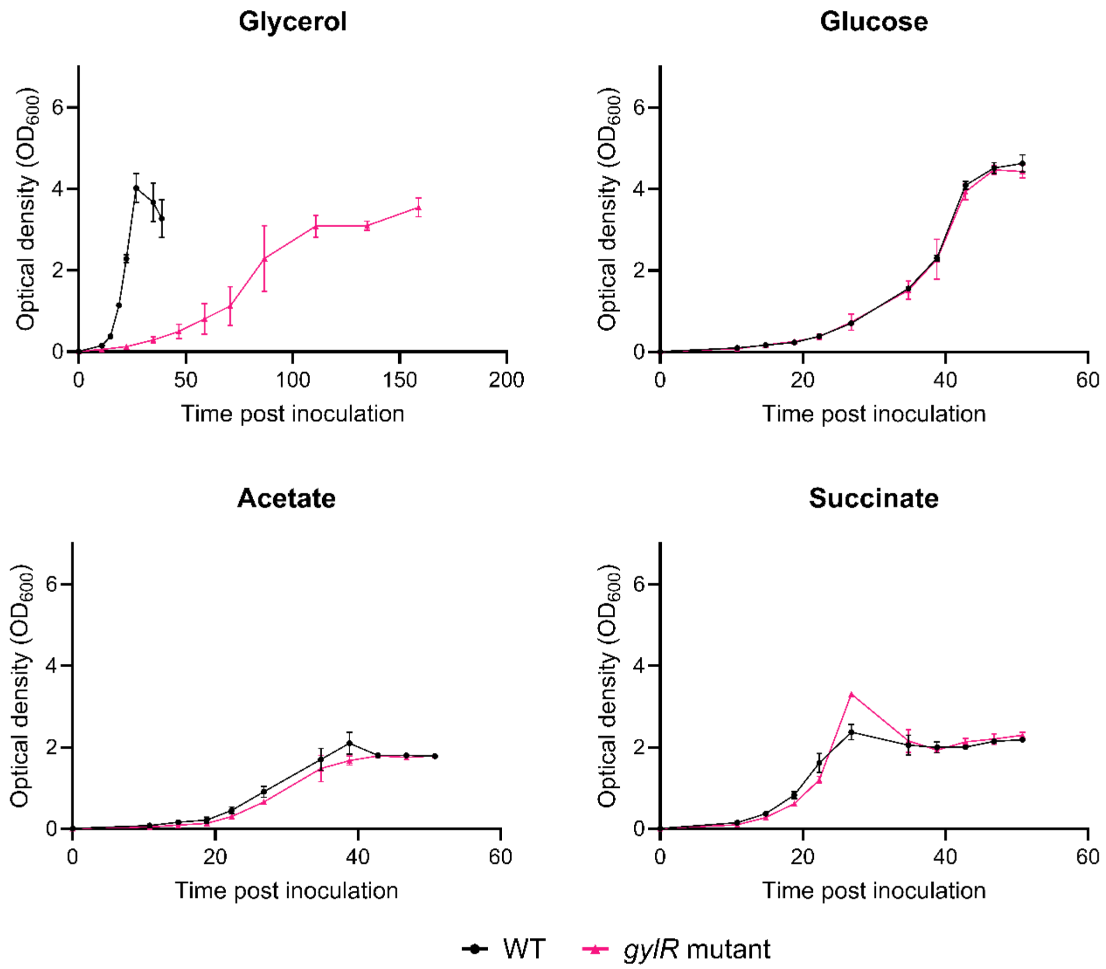


**Figure S2. The presence of a non-functional GylR does not impact the growth of *M. smegmatis* with alternative organic substrates.** Comparative growth of WT *M. smegmatis* and the *gylR* frameshift mutant grown in minimal media supplemented with 0.2% of one of four carbon sources: glycerol, glucose, acetate or succinate. Error bars demonstrate the standard deviations of three (n=3) biological replicates.

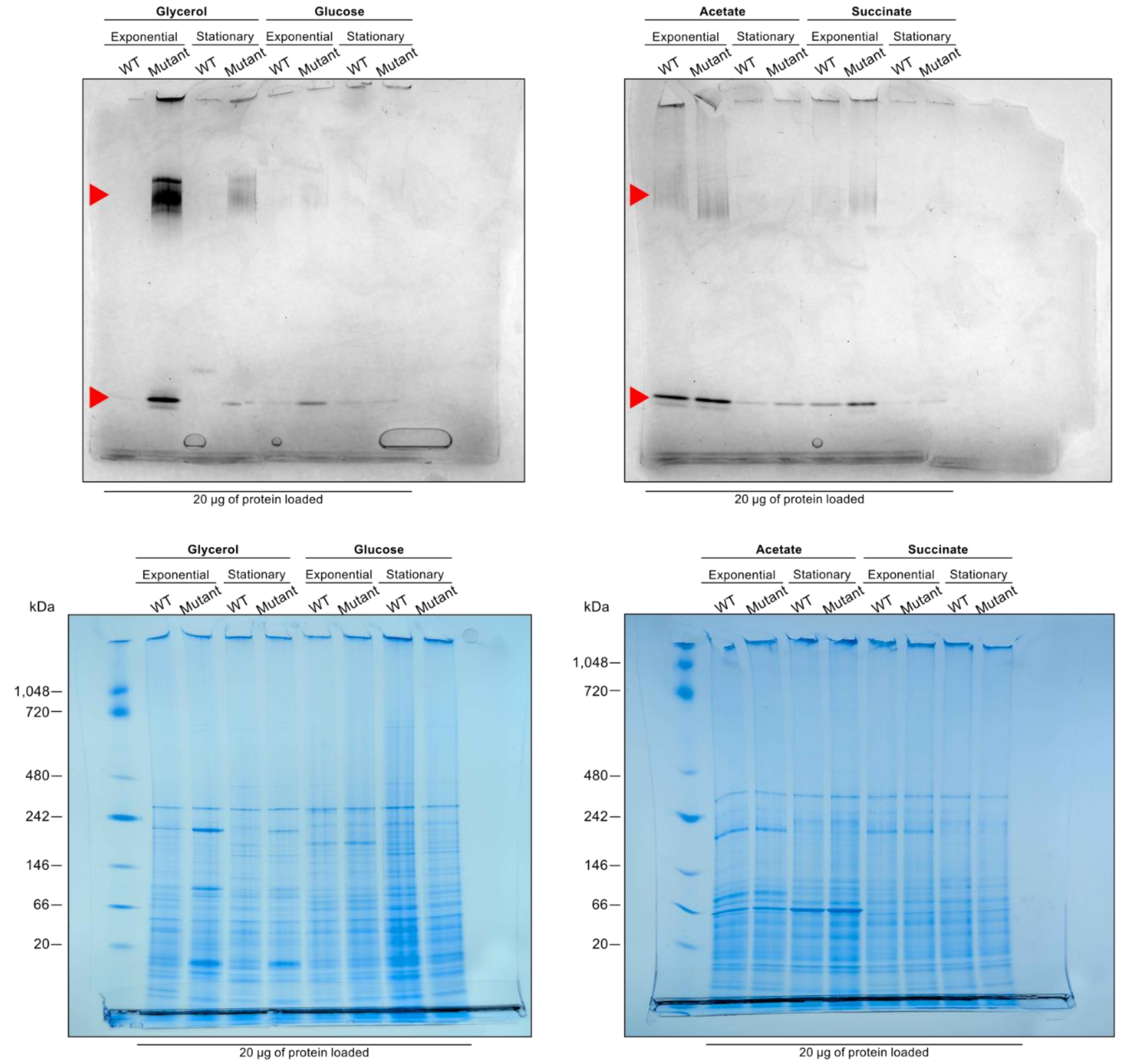


**Figure S3. Native hydrogenase activity staining of WT *M. smegmatis* and *gylR* mutant lysates when grown with different carbon sources.** Native gels depicting hydrogenase activity stained with NBT (top) and Coomassie gel stained with AcquaStain (bottom). Strains were grown with either 0.2% glycerol, glucose, acetate or succinate, with cultures harvested at exponential phase ($OD_{600}$ = 1.3-1.5 for growth with glycerol, $OD_{600}$ = 1.45 for growth with glucose, $OD_{600}$ = 1.2 for growth with succinate, and $OD_{600}$ = 0.9 for growth with acetate) and stationary phase ($OD_{max}$ + 1 day). Red arrows indicate oligomeric Huc staining (upper) and dimeric Huc staining (lower).

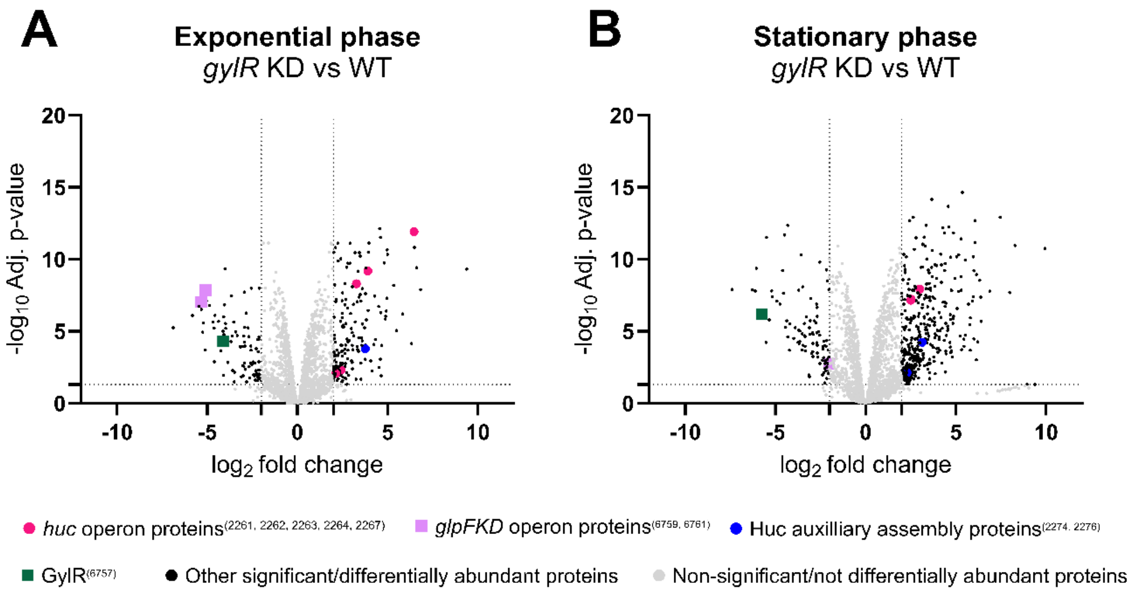


**Figure S4: Comparative proteomic analysis of a CRISPRi-generated knockdown of *gylR* and WT *M. smegmatis*.** Comparative shotgun proteomics volcano plots demonstrating the differential abundance of proteins in the *gylR* knockdown compared to WT *M. smegmatis* when grown with glycerol as the sole carbon source at exponential phase ($OD_{600}$ = 1.5) **(A)** and carbon-depleted stationary phase ($OD_{max}$ + 1 day) **(B)**.

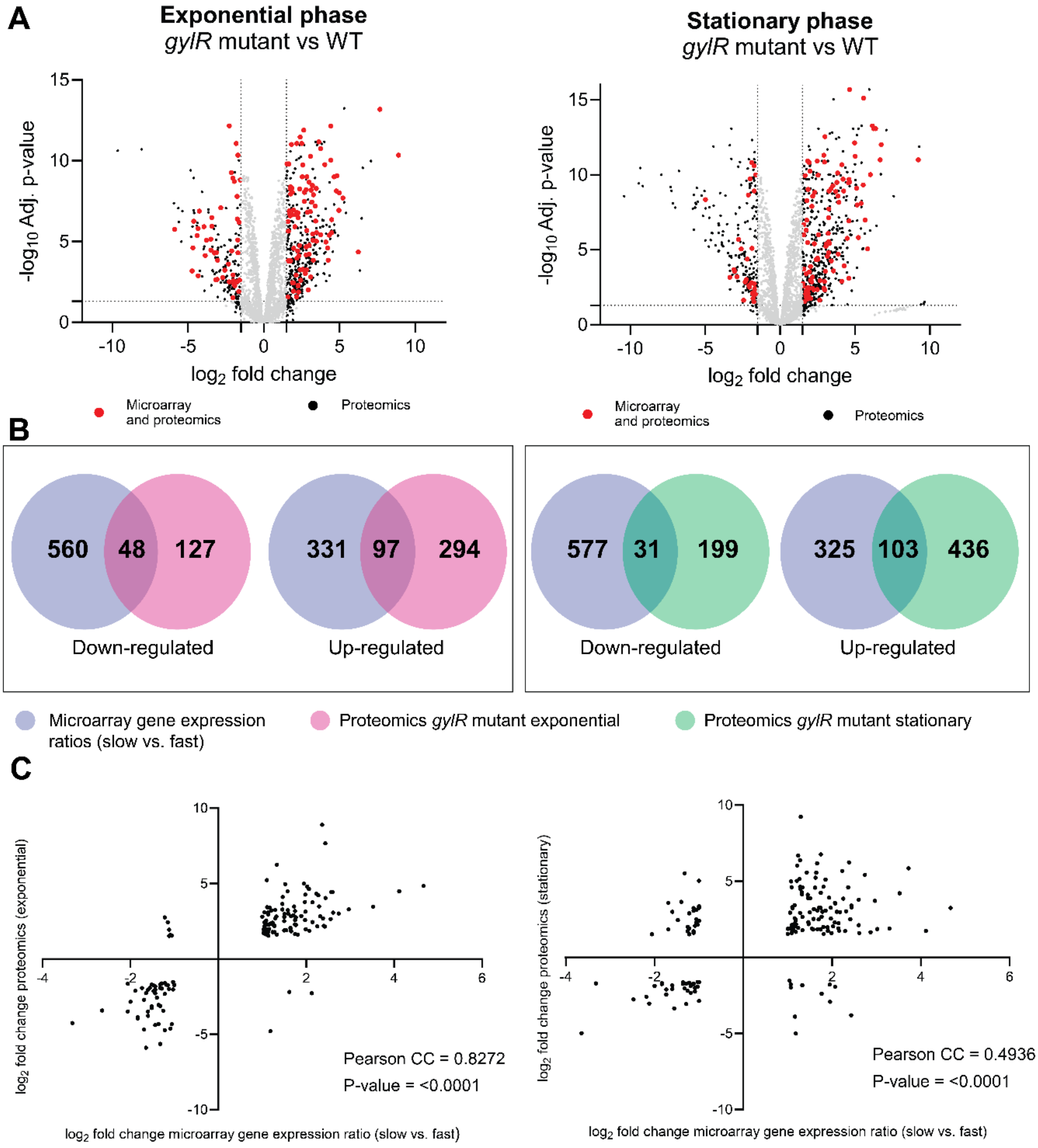


**Figure S5: A comparison of genes differentially expressed in *M. smegmatis* cells during slow vs. fast growth in continuous culture on glycerol and the *gylR* mutant proteomics (A)** Comparative shotgun proteomics volcano plots demonstrating the differential abundance of proteins in the *gylR* mutant compared to WT *M. smegmatis* when grown with glycerol as the sole carbon source at exponential phase ($OD_{600}$ = 1.5) and carbon-depleted stationary phase ($OD_{max}$ + 1 day). Proteins with statistically significant differences in abundance ($p < 0.05$, $log_2$ FC ≥ 1.5, $log_2$ FC ≤ -1.5) are represented by the dark-coloured dots, and those that are not significant are in grey. Red dots indicate proteins that were also found to be up- or down-regulated in microarray data of differentially expressed genes during slow vs. fast growth in continuous culture on glycerol . **(B)** Venn diagrams showing the number of genes significantly up- and down-regulated during slow vs fast growth in continuous culture on glycerol (from microarray data) (blue circle) (39) and proteins with differential abundance in the *gylR* mutant at exponential phase (pink circle) or stationary phase (proteomics data) (green circle). **(C)**

Scatter plots comparing changes in protein abundance in the *gylR* mutant with gene expression-level changes reported in the continuous-culture dataset (39). Only genes/proteins differentially expressed in both datasets are shown. A stronger correlation is observed during exponential growth than in the stationary phase, indicating that the proteomic effects of *gylR* loss more closely mirror transcriptional responses to low glycerol availability during active growth.

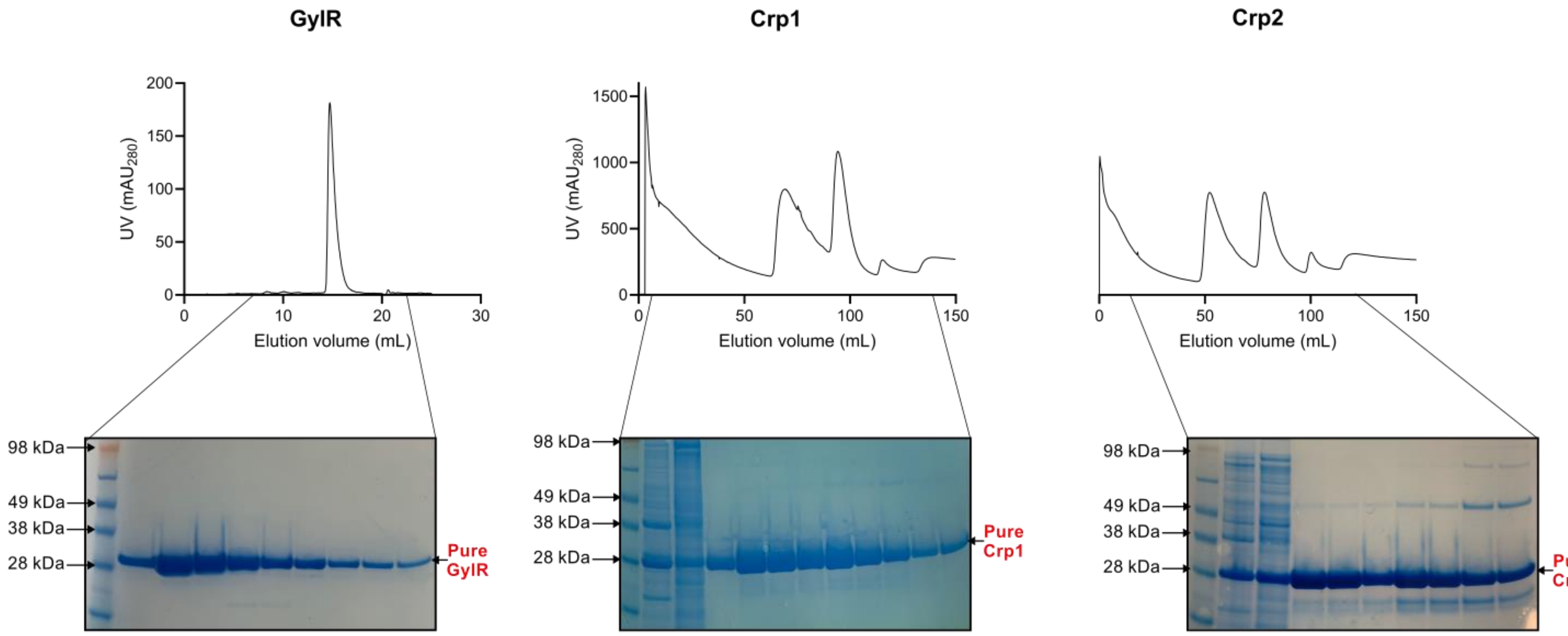


**Figure S6. Purification of recombinantly-expressed GylR, Crp1 and Crp2.** Upper panel: Chromatograms depicting elution profile of GylR (SEC), Crp1 (affinity chromatography) and Crp2 (affinity chromatography). Lower panel: SDS-PAGE gels containing the fractions corresponding to the chromatogram peaks.

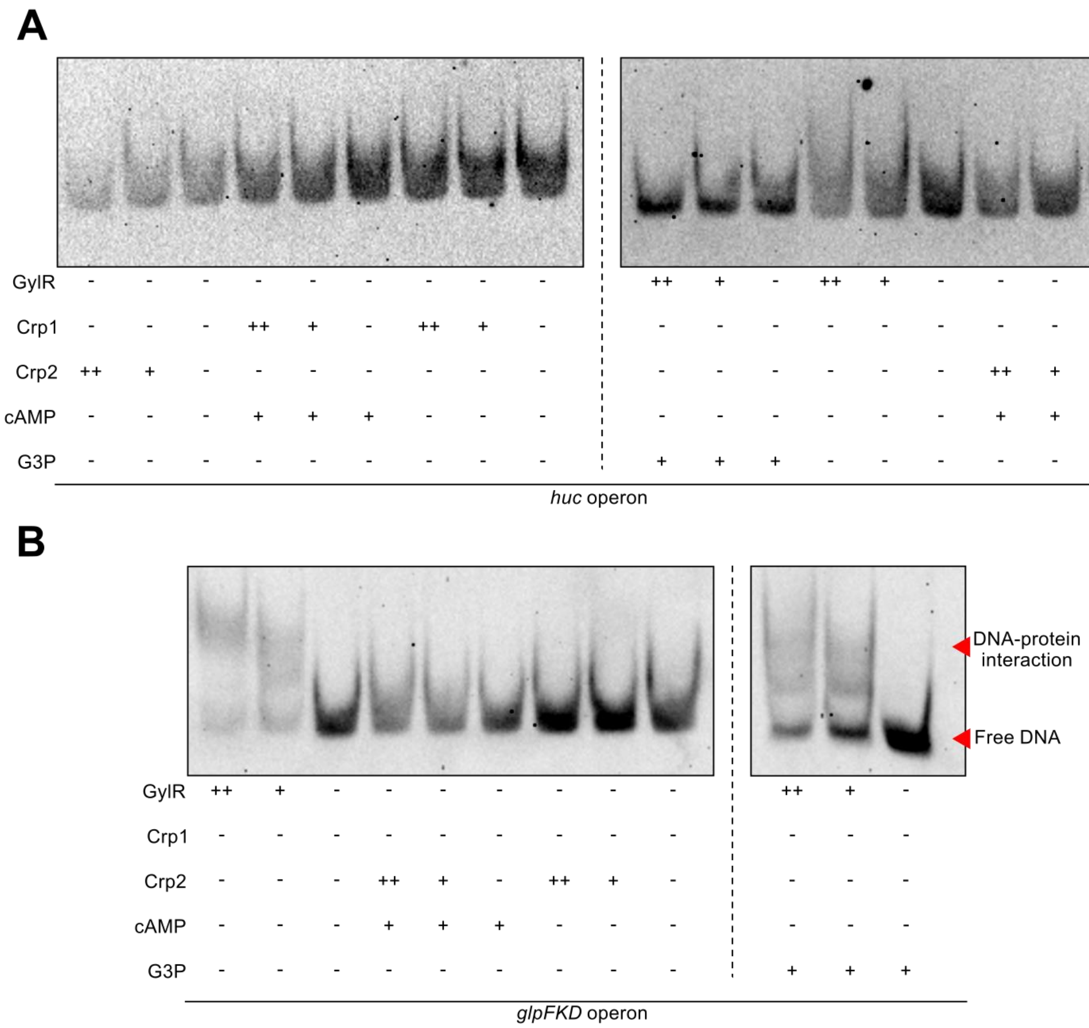


**Figure S7. Electrophoretic Mobility Shift Assays investigating the binding of GylR and CRP to the *huc* promoter region.** Electrophoretic mobility shift assays depicting the binding of GylR and CRP homologues Crp1 and Crp2 to the *huc* **(A)** and *glpFKD* **(B)** operon promoter regions (13, 24). GylR and Crp1/2 were added at a concentration of either 3 µM (++) or 1 µM (+), with cAMP and G3P added to potentially stimulate protein binding at respective concentrations of 50 mM. The binding of GylR to the *glpFKD* promoter was used as a positive control, with DNA-protein interaction indicated by the upward shift in band molecular weight as protein concentration increases.
